# The VINE complex is a VPS9-domain GEF-containing SNX-BAR coat involved in endosomal sorting

**DOI:** 10.1101/2021.11.29.470412

**Authors:** Shawn P. Shortill, Mia S. Frier, Michael Davey, Elizabeth Conibear

## Abstract

Membrane trafficking pathways perform important roles in establishing and maintaining the endolysosomal network. Retrograde protein sorting from the endosome is promoted by conserved SNX-BAR-containing coat complexes including retromer which enrich cargo at tubular microdomains and generate transport carriers. In metazoans, retromer cooperates with VARP, a conserved VPS9-domain GEF, to direct an endosomal recycling pathway. The function of the yeast VARP homolog Vrl1 has been overlooked due an inactivating mutation in commonly studied strains. Here, we demonstrate that Vrl1 has features of a SNX-BAR coat protein and forms an obligate complex with Vin1, the paralog of the retromer SNX-BAR protein Vps5. Unique features in the Vin1 N-terminus allow Vrl1 to distinguish it from Vps5, thereby forming what we have named the VINE complex. VINE occupies endosomal tubules and promotes the delivery of a conserved mannose 6-phosphate receptor-like protein to the vacuolar membrane. In addition to sorting functions, membrane recruitment by Vin1 is essential for Vrl1 GEF activity, suggesting that VINE is a multifunctional coat complex that regulates trafficking and signaling events at the endosome.

## Introduction

Transport of proteins and lipids at the endosome requires the concerted action of peripheral cargo-sorting complexes, Rab-family GTPases, membrane tethering complexes and soluble *N*-ethylmaleimide-sensitive factor attachment protein receptor (SNARE) proteins (Barr and Lambright, 2010; Burd and Cullen, 2014; Jahn and Scheller, 2006; Numrich and Ungermann, 2014; Pfeffer, 2017; Stenmark, 2009; van Weering et al., 2010). Guanine nucleotide exchange factors (GEFs) belonging to the conserved VPS9 family are important regulators of endosomal function that activate endosomal Rab5-like GTPases (Carney et al., 2006; Delprato and Lambright, 2007). In yeast, the VPS9-domain GEFs Muk1 and Vps9 stimulate the endosomal Rabs Vps21, Ypt52 and Ypt53 to perform downstream functions including activating phosphoinositide 3-kinase (PI3K) to produce the anionic lipid species phosphatidylinositol 3-phosphate (PI3P; Christoforidis et al., 1999; Hama et al., 1999; Paulsel et al., 2013; Peplowska et al., 2007; Singer-Krüger et al., 1994). Together, PI3P and endosomal Rabs are important determinants of endosomal identity and are responsible for recruiting effectors, including coat proteins and vesicle tethers, to the endosomal membrane.

Sorting nexins (SNXs) are a conserved family of proteins that perform direct roles in endosomal trafficking by binding to transmembrane cargo proteins and enriching them into sorting domains (Carlton et al., 2005; Hong, 2019). SNXs localize to the endosome through conserved Phox homology (PX) domains that typically recognize PI3P (Cheever et al., 2001; Xu et al., 2001). A sub-family of SNXs known as SNX-BARs additionally contain a Bin/Amphiphysin/RVS (BAR) domain that mediates dimerization with other BAR domain-containing proteins and imparts membrane binding/deforming properties (Frost et al., 2009; van Weering et al., 2012, 2010). These SNX-BAR dimers, which are capable of deforming cargo-rich membranes into sorting tubules, are emerging as important regulators of protein transport. The seven SNX-BAR proteins present in yeast include the conserved retromer subunits Vps5 and Vps17 (Horazdovsky et al., 1997), the SNX8 homolog Mvp1 (Suzuki et al., 2021), the SNX4 homolog Snx4 and its partners Snx41 and Atg20 (SNX7 and SNX30 in humans, respectively; Hettema et al., 2003) and the Vps5 paralog Ykr078w. Retromer and Snx4 complexes promote cargo sorting from the endosome and vacuole (Arlt et al., 2015; Suzuki and Emr, 2018), while Mvp1 appears to function only at the endosome (Suzuki et al., 2021). There is no known function for Ykr078w, which lacks a clear subcellular localization.

The best characterized of these SNX-BAR-containing complexes is the heteropentameric retromer complex, which is composed of the Vps26-Vps35-Vps29 trimer and the Vps5-Vps17 SNX-BAR dimer (Seaman et al., 1998). Retromer localizes to PI3P-rich membranes where it promotes the retrograde sorting of cargo including the well-characterized carboxypeptidase Y (CPY) receptor Vps10 (Bean et al., 2015; Burda et al., 2002; Seaman et al., 1998, 1997). In metazoans, the term retromer refers specifically to the VPS26-VPS35-VPS29 trimer which associates with a variety of adaptor proteins including SNXs to promote cargo sorting (Cullen and Korswagen, 2012; Gallon and Cullen, 2015). Mutations in VPS35 have been linked to neurodegenerative conditions including Parkinson’s disease and Alzheimer’s disease (Mohan and Mellick, 2017; Rahman and Morrison, 2019; Vilariño-Güell et al., 2011; Wen et al., 2011; Zimprich et al., 2011), establishing a connection between endosomal transport machinery and human neurological health.

We previously identified a physical association between yeast retromer and the VPS9-domain GEFs Vps9 and Muk1, whose activity is required to maintain endosomal pools of PI3P for retromer recruitment (Bean et al., 2015). We also identified a novel VPS9-domain protein, Vrl1, which is mutated and non-functional in strains previously used for trafficking studies. Vrl1 can function as the sole VPS9-domain GEF to stimulate production of endosomal PI3P. Notably, the metazoan homolog of Vr1l, VARP, physically associates with retromer to drive an endosome to plasma membrane sorting pathway (Hesketh et al., 2014). Here, we show that Vrl1 is a member of the SNX-BAR protein family and that it specifically binds the Vps5 paralog Ykr078w/Vin1 to form what we now call the VINE complex. Our results suggest that VINE is both a VPS9-domain GEF and a distinct endosomal sorting complex that operates alongside the retromer, Mvp1 and Snx4 pathways.

## Results

### Vrl1 is a predicted PX-BAR protein that interacts with conserved machinery at the endosome

In addition to a conserved VPS9 GEF domain, Vrl1 and VARP share a conserved N-terminus and an ankyrin repeat domain (AnkRD) that are not found in other yeast VPS9-domain GEFs (Figure 1A). Vrl1 also features an unannotated region of ~350 amino acids (aa) downstream of the ankyrin repeats that is not present in VARP. The protein fold recognition program Phyre2 (Kelley et al., 2015) identified a PX-BAR module with very high confidence (98.6%) in this region (aa 737-1089; Figure 1 S1A), and ab initio modeling of this region by the AlphaFold2-powered ColabFold program (Mirdita et al., 2021) predicted a structure with striking similarity to the PX-BAR fold (Figure 1B). Because the predicted Vrl1 PX domain is missing key residues for PI3P binding (Figure 1 S1B), we refer to it as a “PX-like” domain. To our knowledge, Vrl1 is the first VPS9 domain-containing protein with predicted structural homology to the SNX-BAR family.

**Figure 1.**
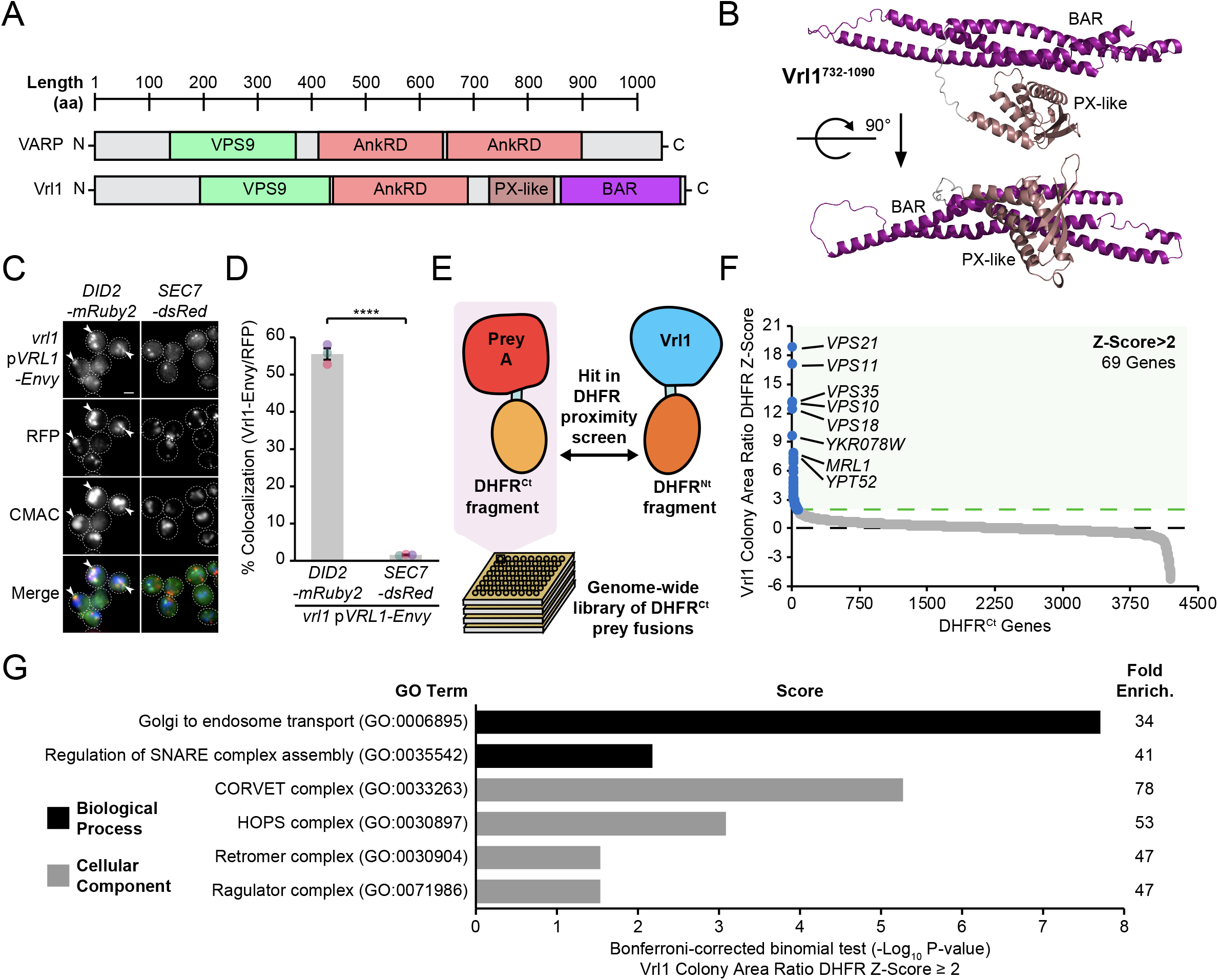
Vrl1 is a predicted PX-BAR protein that interacts with conserved machinery at the endosome. (A) Schematic of Vrl1 and VARP domain architecture. (B) ColabFold predicts the Vrl1 C-terminus has a SNX-BAR-like PX and BAR domain fold. (C) Vrl1-Envy colocalizes with Did2-mRuby2-labeled endosomes, but not with the Sec7-dsRed Golgi marker. (D) Quantification of colocalization as the percentage of Vrl1 puncta over RFP puncta. Two-tailed equal variance *t* test; *n* = 3, cells/strain/replicate ≥ 1,395; **** = P < 0.0001. (E) Schematic of DHFR proximity screen methodology. (F) Z-score distribution of the ratio of colony areas from genome-wide DHFR screens of full length and N-terminal Vrl1 baits. (G) Gene Ontology (GO) functional enrichment analysis of Vrl1 DHFR interactors (Z-score > 2; http://geneontology.org). GO terms of the most specific hierarchical subclass with a fold enrichment value > 25 are presented as the negative base 10 log of the associated P-value from a Bonferroni-corrected binomial test of significance. Scale bars, 2 µm. Error bars report standard error of the mean (SEM). Enrich., enrichment. aa, amino acids.

We found that Vrl1, when C-terminally tagged with the bright GFP variant Envy, was present at perivacuolar puncta that colocalize with the endosomal marker Did2-mRuby2, but not the Golgi protein Sec7-dsRed (56% and 2%, respectively, P < 0.0001; Figure 1C and 1D). These observations indicate that unlike other yeast VPS9-domain GEFs (Paulsel et al., 2013), Vrl1 constitutively localizes to endosomes. To identify endosomal partners of Vrl1, we performed a protein fragment complementation assay (PCA) based on a drug-resistant variant of the dihydrofolate reductase (DHFR) enzyme (Figure 1E; Michnick et al., 2010; Tarassov et al., 2008). Proximity between two proteins that are fused to complementary DHFR fragments reconstitutes enzyme activity and confers resistance to the inhibitor methotrexate. Full length Vrl1, and a cytosolic fragment of Vrl1 lacking the AnkRD, PX-like and BAR domains (Figure 1 S2A), were expressed as DHFR^Nt^ fusions under the control of the constitutive *ADH1* promoter (*ADH1pr)*. Z-scores were generated from the colony area ratio of full-length vs truncated Vrl1 baits (Figure 1F, Table S1). This identified the endosomal Rab GTPases Vps21 (Z = 18.7) and Ypt52 (Z = 7.6), and other conserved endosomal proteins including the retromer subunit Vps35 (Z = 13), the hydrolase receptors Vps10 (Z = 12.9) and Mrl1 (Z = 7.7), and components of the Class C Core complex Vps11 (Z = 16.9) and Vps18 (Z = 12.2; Figure 1F). Functional enrichment analysis of Vrl1 interactors (Z > 2) highlighted relationships with other subunits of endosomal complexes including retromer and the CORVET complex (Figure 1G, Table S2; Ashburner et al., 2000; Gene Ontology Consortium, 2021). These results suggest that Vrl1 is an endosomal SNX-BAR-like protein that contacts both membrane tethering and trafficking machinery.

### Vrl1 and the Vps5 paralog Vin1 form the VINE complex

Our DHFR screen identified a strong link between Vrl1 and the uncharacterized SNX-BAR Ykr078w (Z = 9.5), the paralog of membrane-binding retromer subunit Vps5 (Byrne and Wolfe, 2005; Horazdovsky et al., 1997), which we have named “Vrl1-Interacting Sorting Nexin 1” or Vin1 (Figure 2A). Vin1 has a reported cytosolic distribution (Huh et al., 2003), which is surprising given that its paralog Vps5 localizes to endosomes in a PI3P-dependent manner (Burda et al., 2002) and that Vin1 interacts with PI3P in vitro (Yu and Lemmon, 2001). Increasing *VIN1* expression did not alter this cytosolic distribution pattern (Figure 2 S1A), suggesting that a specific condition or recruitment factor may be missing. Indeed, we found that expression of *VRL1* caused a dramatic redistribution of Vin1-Envy from the cytosol to intracellular puncta (P < 0.0001; Figure 2B and C), and that expressing *VRL1* from the strong *ADH1pr* further increased the number of bright Vin1-Envy puncta (P < 0.0001; Figure 2 S1B and S1C). Deletion of *VIN1* prevented Vrl1-Envy from forming intracellular puncta (P < 0.001; Figure 2B and 2C), suggesting that the localization of Vrl1 and Vin1 is highly interdependent.

**Figure 2.**
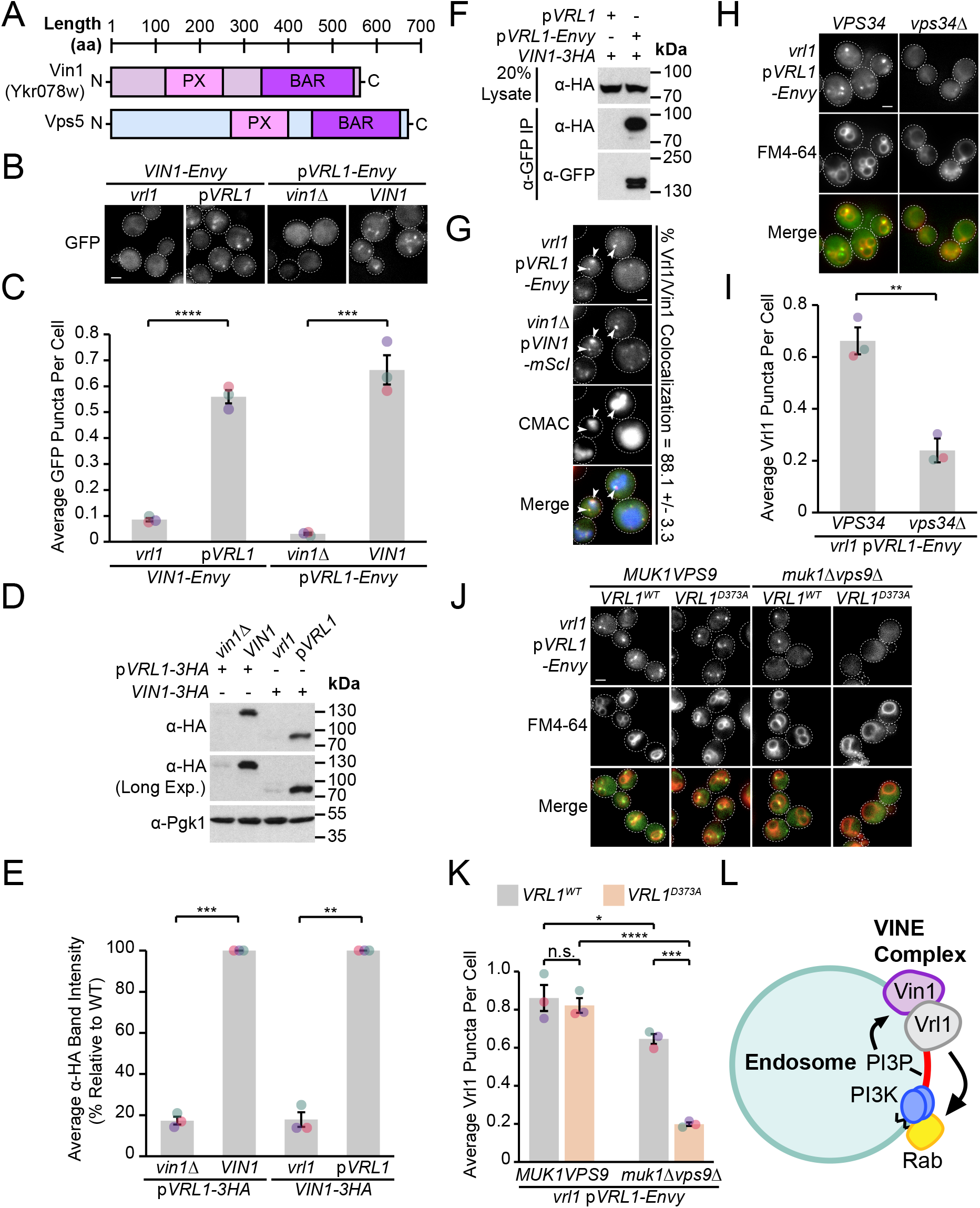
Vrl1 and the Vps5 paralog Vin1 form the VINE complex. (A) Schematic of Ykr078w (Vin1) and its paralog Vps5. (B) Vin1-Envy and Vrl1-Envy require Vrl1 and Vin1, respectively, for localization to puncta. (C) Quantification of Vin1-Envy and Vrl1-Envy puncta per cell. Two tailed equal variance *t* tests; *n* = 3, cells/strain/replicate ≥ 1,879; *** = P < 0.001, **** = P < 0.0001. (D) Vrl1-3HA and Vin1-3HA require Vin1 and Vrl1, respectively, for protein stability by western blot. Pgk1 serves as a loading control. (E) Quantification of Vrl1-3HA and Vin1-3HA levels by densitometry. Two tailed Welch’s *t* tests; *n* = 3, ** = P < 0.01, *** = P < 0.001. (F) Co-immunoprecipitation (CoIP) of Vin1-3HA with Vrl1-Envy suggests stable complex formation. (G) Vrl1-Envy colocalizes with Vin1-mScI at perivacuolar puncta. (H) Vrl1-Envy requires the PI3K catalytic subunit Vps34 for punctate localization. (I) Quantification of Vrl1-Envy puncta per cell. Two-tailed equal variance *t* test; *n* = 3, cells/strain/replicate ≥ 897; ** = P < 0.01. (J) Vrl1-Envy localization in the absence of VPS9-domain GEFs is dependent on Vrl1 activity. (K) Quantification of Vrl1-Envy puncta per cell. One-way ANOVA with Tukey’s multiple comparison test; *n* = 3, cells/strain/replicate ≥ 1,705; not significant, n.s. = P > 0.05, * = P < 0.05, *** = P < 0.001, **** = P < 0.0001. (L) model of the Vin1 and Vrl1-containing VINE complex at endosomes. Scale bars, 2 µm. Error bars report SEM. Exp., Exposure

Vrl1 and Vin1 are also dependent on each other for stability, as the levels of triple hemagglutinin (3HA) tagged Vrl1 and Vin1 were severely reduced in strains lacking *VIN1* or *VRL1* (17% of WT, P < 0.001 and 18% of WT, P < 0.01, respectively; Figure 2D and 2E). We found that Vin1-3HA strongly co-purified with Vrl1-Envy (64% recovery of Vin1; Figure 2F), and Vrl1-Envy and Vin1-mScarletI (-mScI) showed a high degree of colocalization at endosomal puncta (88%; Figure 2G), suggesting that these proteins form a complex.

Since both Vin1 and Vrl1 have predicted PX domains, we wondered if, like other SNX-BARs, they bind PI3P at endosomes. Vrl1 was displaced to the cytosol in a PI3K deletion mutant (*vps34*Δ, P < 0.01; Figure 2H and 2I). Since the Vrl1 PX domain is missing residues that are typically required to bind PI3P, this suggests the PI3P-binding PX domain of Vin1 is important for endosomal recruitment. PI3K is activated by Rab5-like GTPases (Christoforidis et al., 1999), which in turn require VPS9-domain GEFs for their activity (Carney et al., 2006; Delprato and Lambright, 2007). We found that, in a *muk1*Δ*vps9*Δ strain that lacks all other VPS9-domain GEFs, Vrl1 becomes dependent on its own GEF activity for localization (P < 0.001; Figure 2J and 2K), suggesting that it may leverage its ability to stimulate endosomal PI3P production and promote its own membrane recruitment.

Taken together, our results suggest that Vrl1 and Vin1 form a novel complex that localizes to endosomes in a PI3P-dependent manner (Figure 2L). Since neither Vrl1 or Vin1 are stable or capable of membrane localization in the absence of the other, we reason that these proteins primarily exist as members of this complex which we have named the “VPS9 GEF-Interacting Sorting Nexin” or VINE complex.

### Vrl1 interacts with Vin1 primarily via the AnkRD

SNX-BAR proteins interact via an extensive hydrophobic interface between the BAR domains (van Weering et al., 2012). The ColabFold algorithm (Mirdita et al., 2021) predicted that the PX-like and BAR domains of Vrl1 (aa 732-1090) bind to the PX and BAR domains of Vin1 (aa 110-585) to form a canonical SNX-BAR dimer (pTMscore = 0.75; Figure 3A). To assess the accuracy of ColabFold in predicting specific BAR domain pairings, we systematically modelled pairwise homotypic and heterotypic interactions of several yeast SNX-BAR proteins (Figure 3B, Table S3). This accurately predicted the homodimerization of Mvp1 (Suzuki et al., 2021) and the heterodimerization of Vps5/Vps17 (Seaman and Williams, 2002). Neither Vrl1 nor Vin1 were predicted to form homodimers, though unexpectedly Vrl1 was predicted to pair equally well with both Vin1 and its paralog Vps5.

**Figure 3.**
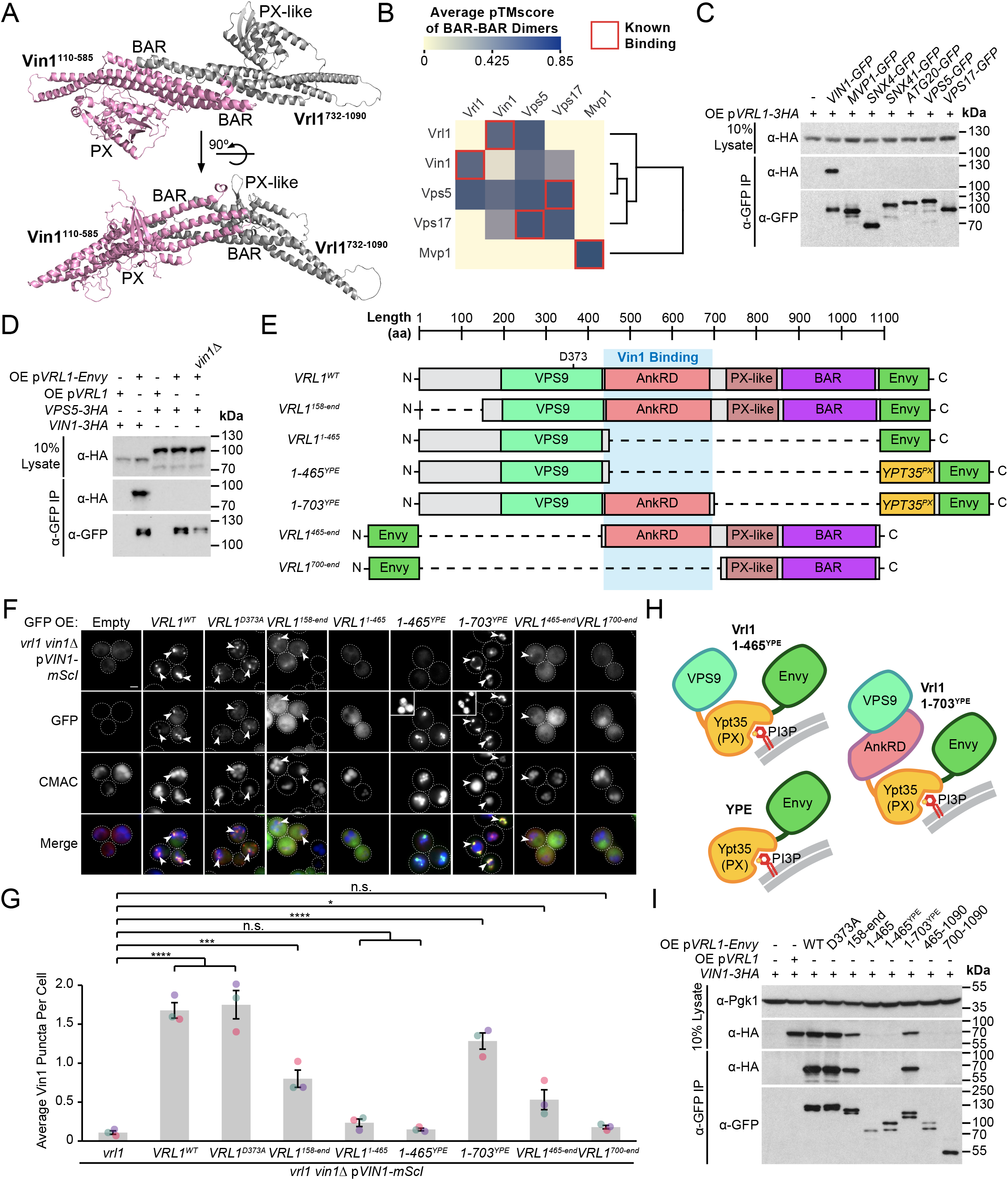
Vrl1 interacts with Vin1 primarily through the AnkRD. (A) ColabFold-predicted physical interaction of Vrl1 and Vin1 BAR domains along the canonical BAR-BAR dimerization interface. pTMscore = 0.75. (B) Matrix of ColabFold-predicted BAR-BAR dimers for selected yeast SNX-BARs. Hierarchical clustering was performed using an uncentered Pearson correlation with average linkage. (C) Vin1 is the only yeast SNX-BAR that interacts with overexpressed Vrl1-3HA by CoIP. (D) Vin1 does not compete with Vps5 for shared binding to Vrl1. (E) Schematic of Envy-tagged Vrl1 truncations and chimeras in (F). (F) The Vrl1 AnkRD is necessary to recruit Vin1-mScI to puncta. Insets are scaled to match other regions in the same channel. (G) Quantification of Vin1-mScI puncta per cell. One-way ANOVA with Dunnett’s multiple comparison test; *n* = 3, cells/strain/replicate ≥ 764; not significant, n.s. = P > 0.05, * = P < 0.05, ** = P < 0.01, *** = P < 0.001, **** = P < 0.0001. (H) Illustration of chimeric Vrl1 fusion proteins that are artificially recruited to the endosomal system by the PX domain of sorting nexin Ypt35. (I) The Vrl1 AnkRD is necessary for physical interaction with Vin1-3HA by CoIP. Pgk1 serves as a loading control. Scale bars, 2 µm. Error bars report SEM. OE, overexpressed. YPE, Ypt35(PX)-Envy.

We wondered if Vrl1 could also partner with Vps5 to form a novel retromer-like complex. First, we tested all known yeast SNX-BAR proteins for their ability to bind Vrl1 (Figure 3C) and found that Vin1 is the only yeast SNX-BAR that binds Vrl1 at a detectable level in cell lysates. Vrl1 also failed to bind Vps5 when *VIN1* was deleted (Figure 3D), and we found no evidence that Vrl1 partners with other retromer subunits (Figure 3 S1A), but this assay could fail to detect weak or transient interactions. Using functional readouts, we found that Vrl1 was unable to promote the endosomal localization of Vps10 (Figure 3 S1B) or Vps35 (Figure 3 S1C and S1D) in strains lacking Vps5 and/or Vps17, indicating that Vrl1 does not functionally pair with retromer SNX-BARs and that Vin1/Vrl1 cannot replace Vps5/17 to form a retromer-like complex. These results further suggest that Vrl1 has strong isoform specificity and that interactions beyond the BAR-BAR interface could be responsible for its specific recognition of Vin1.

To identify regions critical for Vrl1-Vin1 binding, we quantified the membrane recruitment of Vin1-mScarletI in cells expressing a series of Envy-tagged Vrl1 fragments (Figure 3E and 3F). We found that the GEF-deficient mutant (Vrl1^D373A^; Bean et al., 2015) and the N-terminal truncation (Vrl1^158-end^) significantly recruited Vin1 (P < 0.0001 and P < 0.001, respectively; Figure 3G) despite the weaker punctate localization of the Vrl1^158-end^ construct relative to WT, which could explain its reduced recruitment of Vin1. Deletion of C-terminal sequences (eg Vrl1^1-465^) blocked the membrane localization of both Vrl1 and Vin1. To test the role of the Vrl1 PX-like and BAR domains, we replaced this region with a localization module consisting of the PI3P-binding PX domain of Ypt35 fused to Envy which we refer to as “YPE” (Figure 3H). Strikingly, the resulting Vrl1(1-703)^YPE^ chimera strongly recruited Vin1 to puncta (P < 0.0001; Figure 3G), suggesting that BAR-BAR interactions are dispensable for Vin1 recruitment. A further truncation that removed the AnkRD to create Vrl1(1-465)^YPE^ localized to endosomes yet failed to recruit Vin1 (Figure 3F and 3G), indicating that the AnkRD contains a potent interacting interface for Vin1. In support of this idea, the Vrl1^465-end^ fragment which contains the AnkRD, PX-like and BAR domains weakly localized and recruited a small but significant amount of Vin1 (P < 0.05) while the Vrl1^700-end^ fragment containing only the PX-like and BAR domains did not localize or recruit Vin1 (Figure 3F and Figure 3G).

We then tested the Vrl1 truncation series (Figure 3E) for the ability to CoIP Vin1-3HA (Figure 3I). We found that, except for Vrl1^465-end^ which weakly recruited Vin1 to membranes, all constructs that were able to recruit Vin1 physically bound it by CoIP. Taken together, the Vin1 recruitment/binding assays and structural predictions suggest that the VINE complex assembles primarily through an interaction between Vin1 and the Vrl1 AnkRD, while a secondary interaction between the Vin1 and Vrl1 BAR domains may occur at the endosomal membrane.

### The Vrl1 AnkRD recognizes a small region of the Vin1 N-terminus

AnkRD interactions may explain how Vrl1 discriminates between Vin1 and its paralog Vps5 (Figure 3C and 3D). Vin1 and Vps5 have extended, divergent N-terminal sequences preceding their respective PX domains (Vin1^1-116^ and Vps5^1-276^; Figure 4A). When we expressed the N-terminal regions of Vin1 and Vps5 fused to mScarletI (Figure 4B), we observed strong recruitment of the Vin1 N-terminus by the Vrl1(1-703)^YPE^ chimera, but not the YPE module alone (P < 0.0001; Figure 4C). In contrast, we detected no recruitment of the Vps5 N-terminus by any of our tested constructs suggesting that the N-terminal regions of the paralogous SNX-BARs dictate specificity for Vrl1.

**Figure 4.**
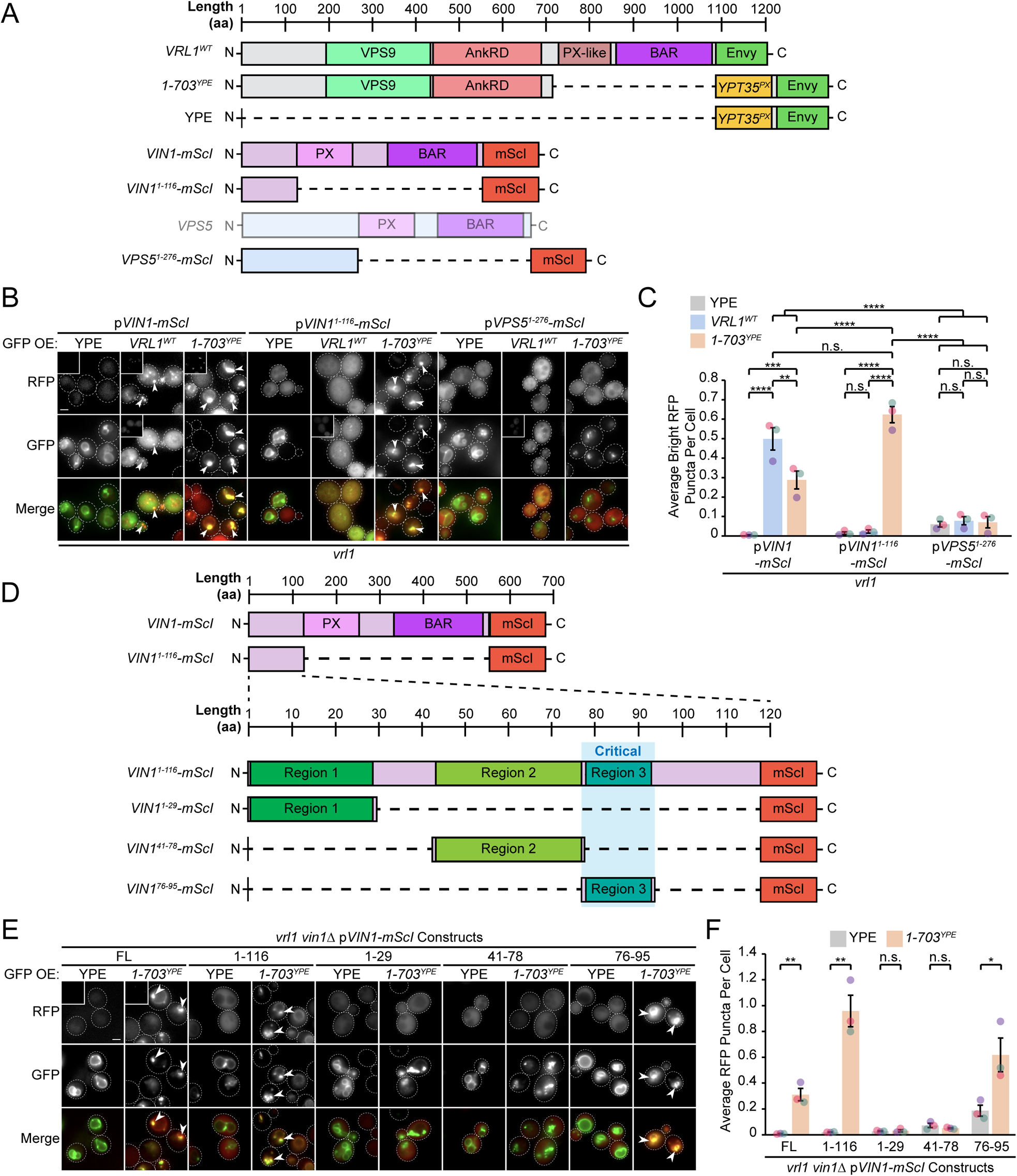
The Vrl1 AnkRD recognizes a small region of the Vin1 N-terminus. (A) Schematic of constructs used in (B, C) (B) The AnkRD-containing Vrl1-YPE chimera recruits the N-terminus of Vin1, but not Vps5. Insets are scaled to match other regions in the same channel. (C) Quantification of RFP puncta per cell. One-way ANOVA with Tukey’s multiple comparison test; *n* = 3, cells/strain/replicate ≥ 902; not significant, n.s. = P > 0.05, ** = P < 0.01, *** = P < 0.001, **** = P < 0.0001. (D) Schematic of Vin1 N-terminal fragments used to map the Vrl1 recruitment site. (E) The AnkRD-containing Vrl1-YPE chimera recruits a small fragment of the Vin1 N-terminus. Insets are scaled to match other regions in the same channel. (F) Quantification of Vin1-mScI puncta per cell. Two-tailed equal variance *t* tests; *n* = 3, cells/strain/replicate ≥ 294; not significant, n.s. = P > 0.05, * = P < 0.05, ** = P < 0.01. Scale bars, 2 µm. Error bars report SEM. OE, overexpressed. FL, full length.

We noticed that WT Vrl1-Envy recruited WT Vin1-mScI to colocalizing puncta, (Figure 4B) but was unable to recruit the Vin1 N-terminus. The endogenous, untagged Vin1 may outcompete the Vin1 N-terminal fragment for recruitment by Vrl1, however this could not be tested directly because WT Vrl1 failed to localize when the Vin1 N-terminus was expressed in a *vin1*Δ strain (Figure 4 S1A and S1B). This observation indicates that the Vin1 PX and BAR domains also contribute to VINE assembly and membrane recruitment.

We generated an alignment from fungal ohnologs of Vps5 and Vin1 (Figure 4 S2A; Byrne and Wolfe, 2005) that identified three conserved regions in the Vin1 N-terminus (Figure 4D). When each of these fragments was fused to mScarletI (Figure 4E), only region 3 (Vin1 aa 76-95) was recruited to puncta by Vrl1(1-703)^YPE^ in a *vin1*Δ strain (P < 0.05; Figure 4F). This suggests that the Vrl1 AnkRD distinguishes Vin1 from Vps5 through a short sequence in the unstructured Vin1 N-terminus.

### Vin1 regulates Vrl1 GEF activity via membrane localization

We used different functional readouts to test if Vin1 is required for Vrl1 GEF activity. Disruption of the other VPS9-domain GEF proteins results in a severe temperature sensitivity phenotype and loss of endosomal PI3P (Paulsel et al., 2013; Singer-Krüger et al., 1994) that is rescued by Vrl1 in an activity-dependent manner (Bean et al., 2015). We found that deletion of *VIN1,* but not *VPS5*, prevented Vrl1 from rescuing the temperature sensitivity of the *muk1Δvps9Δ* strain (Figure 5A). We also found that localization of a fluorescent PI3P biosensor (Figure 5B) was restricted to the vacuolar membrane in *muk1Δvps9Δvin1Δ* cells expressing Vrl1 (Figure 5C), suggesting that VINE promotes the synthesis of endosomal PI3P only when fully assembled.

**Figure 5.**
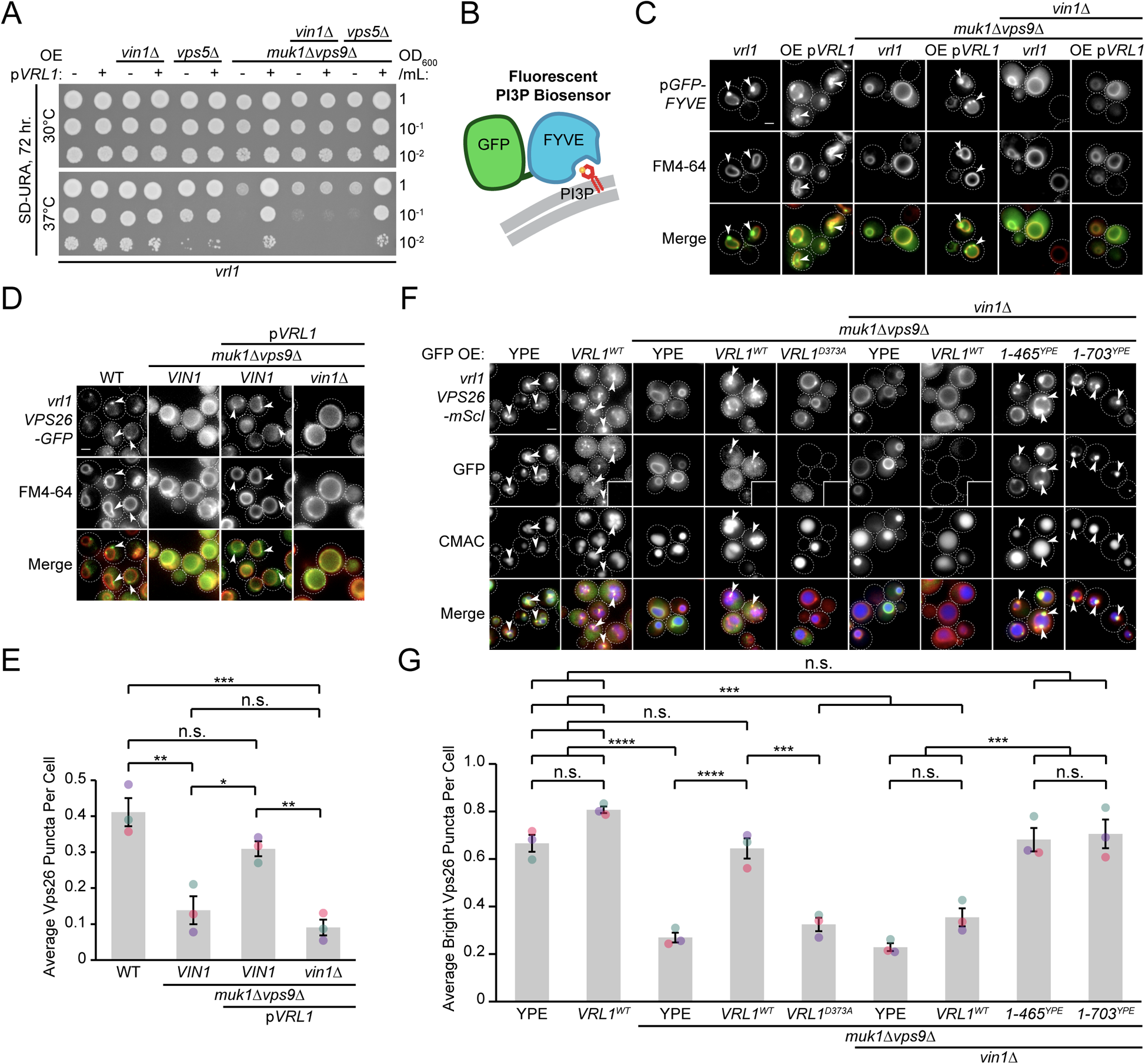
Vin1 controls Vrl1 GEF activity via membrane localization. (A) Deletion of *VIN1*, but not *VPS5*, prevents Vrl1 from rescuing the temperature sensitivity of a strain lacking other VPS9-domain GEFs. (B) Schematic of PI3P-binding fluorescent biosensor. (C) Deletion of *VIN1* prevents Vrl1 from stimulating endosomal PI3P production in a strain lacking other VPS9-domain GEFs. (D) Deletion of *VIN1* prevents Vrl1 from rescuing Vps26-GFP localization in a strain lacking other VPS9-domain GEFs. (E) Quantification of Vps26-GFP puncta per cell. One-way ANOVA with Tukey’s multiple comparison test; *n* = 3, cells/strain/replicate ≥ 1,503; not significant, n.s. = P > 0.05, * = P < 0.05, ** = P < 0.01, *** = P < 0.001. (F) Vin1 is dispensable for Vrl1 activity when fragments containing the N-terminus and VPS9 domain are artificially recruited by a YPE endosomal anchor. Insets are scaled to match other regions in the same channel. (G) Quantification of Vps26-mScI puncta per cell. One-way ANOVA with Tukey’s multiple comparison test; *n* = 3, cells/strain/replicate ≥ 750; not significant, n.s. = P > 0.05, *** = P < 0.001, **** = P < 0.0001. Scale bars, 2 µm. Error bars report SEM. OE, overexpressed.

We previously found that Vrl1 recovers the PI3P-dependent endosomal localization of retromer in a *muk1*Δ*vps9*Δ strain (Bean et al., 2015). By quantifying the localization of the endogenously tagged retromer subunit Vps26-GFP (Figure 5D), we reproduced this finding and found that deletion of *VIN1* blocked rescue (P < 0.01; Figure 5E). We next tested if Vin1 was still required for Vrl1 activity when the PX-like and BAR domains of Vrl1 were replaced by the YPE endosomal anchor (Figure 3H) using Vps26-mScarletI localization as a readout (Figure 5F). We observed that Vrl1(1-465)^YPE^ and Vrl1(1-703)^YPE^, but not WT Vrl1, fully rescued the endosomal localization of Vps26-mScarletI in a *muk1*Δ*vps9*Δ*vin1Δ* strain (P < 0.001; Figure 5G). These results suggest that Vin1 regulates the activity of Vrl1 by promoting its localization to endosomes.

### The VINE complex exhibits characteristics of a SNX-BAR membrane sorting complex

We wondered if the VINE complex occupies endosomal membrane tubules as other SNX-BAR coat complexes do (Suzuki et al., 2021; van Weering et al., 2012; Zhang et al., 2021). To test this, we co-overexpressed Vrl1 and GFP-Vin1 on the strong *ADH1* and *NOP1* promoters, respectively, and acquired images of GFP-Vin1 at 100ms intervals (Figure 6A). When compared to the endosomal marker Did2-mRuby2, we could observe GFP-Vin1 in tubular structures that eventually underwent scission and separated from the endosome (Figure 6A and 6B).

**Figure 6.**
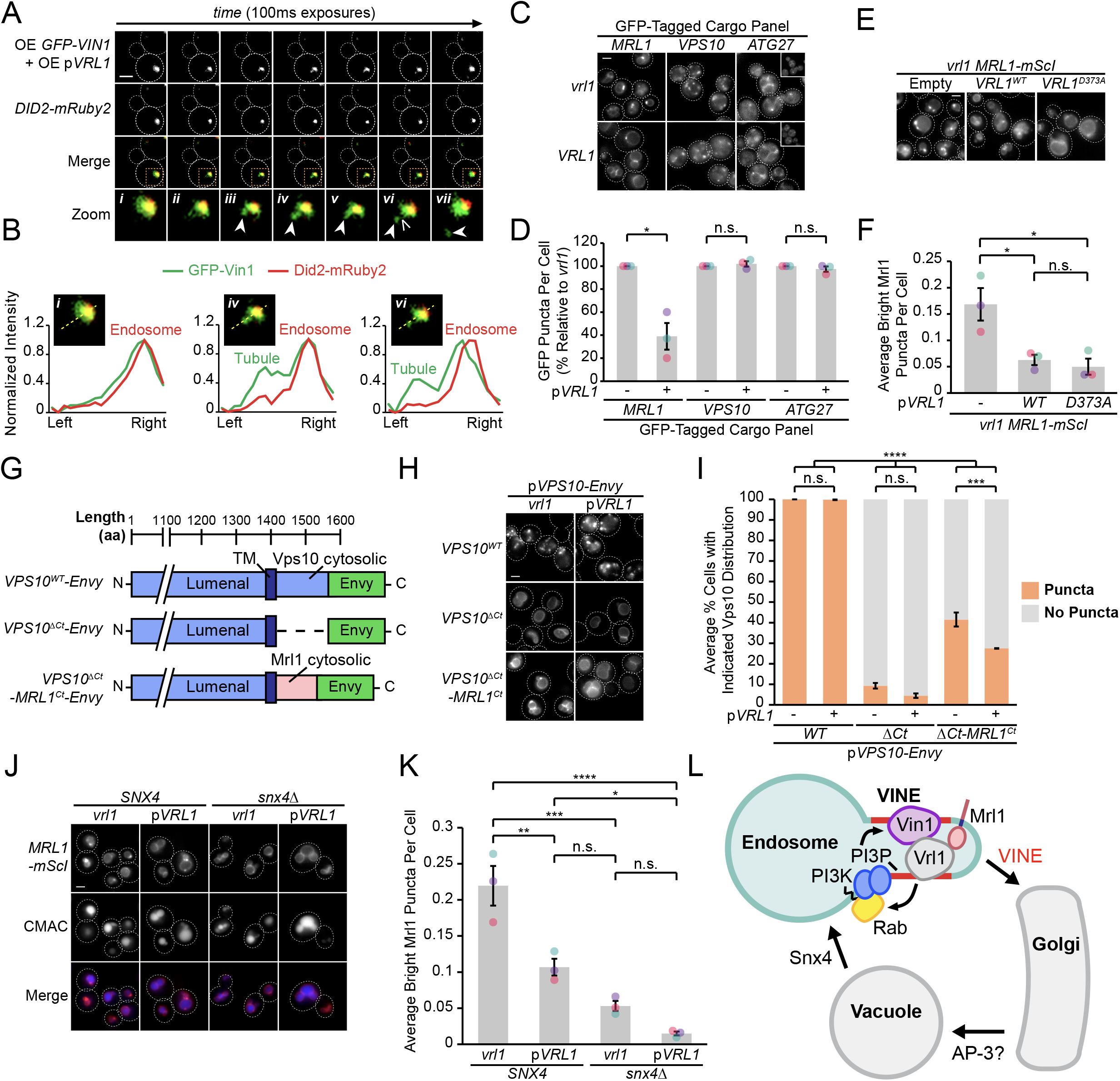
The VINE complex exhibits characteristics of a membrane sorting complex. (A) Time-lapse imaging of cells co-overexpressing GFP-Vin1 and Vrl1 show tubules emanating from Did2-labeled endosomes. Images were uniformly enlarged using a bicubic expansion function to show detail. Solid arrowheads mark a tubule, open arrowhead marks a scission event. (B) Normalized intensity line scan analysis performed on images from (A) along the yellow dotted line. (C) Punctate localization of GFP-tagged Mrl1, but not other endosomal recycling cargo, is decreased in cells expressing *VRL1*. (D) Quantification of GFP-tagged puncta in WT and *vrl1* strains. Two tailed Welch’s *t* tests; *n* = 3, cells/strain/replicate ≥ 902; not significant, n.s. = P > 0.05, * = P < 0.05. (E) Mutation of the D373 residue required for VPS9 GEF activity does not prevent Vrl1 from redistributing Mrl1. (F) Quantification of Mrl1-mScI puncta per cell. One-way ANOVA with Tukey’s multiple comparison test; *n* = 3, cells/strain/replicate ≥ 1,788; not significant, n.s. = P > 0.05, * = P < 0.05. (G) Schematic of Vps10 cytosolic tail mutant and Mrl1 cytosolic tail chimera tested for VINE-mediated sorting in (H, I). (H) The Mrl1 cytosolic tail is sufficient to confer VINE-mediated redistribution. (I) Percent of cells showing punctate localization of indicated GFP-tagged constructs. Blind scoring of GFP signal was conducted manually. One-way ANOVA with Tukey’s multiple comparison test; *n* = 3, cells/strain/replicate ≥ 237; not significant, n.s. = P > 0.05, *** = P < 0.001, **** = P < 0.0001. (J) Mrl1-mScI puncta are reduced in a *snx4Δ* strain. (K) Quantification of Mrl1-mScI puncta per cell. One-way ANOVA with Tukey’s multiple comparison test; *n* = 3, cells/strain/replicate ≥ 1,036; not significant, n.s. = P > 0.05, * = P < 0.05, ** = P < 0.01, *** = P < 0.001, **** = P < 0.0001. (L) Model for VINE-mediated sorting of Mrl1. VINE recognizes Mrl1 through its cytosolic tail and transports it from the endosome to the Golgi, where it is subsequently delivered to the vacuolar membrane. Scale bars, 2 µm. Error bars report SEM. OE, overexpressed. TM, transmembrane.

The budding of VINE-coated endosomal tubules suggests the VINE complex transports cargo proteins from this organelle. We examined the localization of several candidates in the presence and absence of Vrl1 (Figure 6C), including the mannose 6-phosphate receptor (MPR) homolog Mrl1, which had a strong DHFR interaction score with Vrl1 (Z = 7.7; Figure 1G), and two proteins that require other SNX-BARs for their transport (Seaman et al., 1998, 1997; Suzuki and Emr, 2018; Suzuki et al., 2021). We found that *VRL1* expression had no significant effect on the localization of the retromer cargo Vps10 or the Snx4 cargo Atg27 but caused a significant decrease in the number of bright Mrl1-GFP puncta per cell (61% decrease relative to *vrl1*, P < 0.05; Figure 6D). Vrl1 GEF activity was not required for this effect (P < 0.05; Figure 6E and 6F), indicating that Vrl1 does not alter Mrl1 distribution by influencing the local activity of endosomal Rabs. Correction of the *vrl1* frameshift mutation using CRISPR-Cas9 gene editing technology restored the punctate localization of Vin1 (Figure 6 S1A) and caused a similar change in Mrl1 localization (P < 0.01; Figure 6 S1B and S1C). This Vrl1-dependent change in Mrl1 distribution strongly suggests that the VINE complex regulates the Mrl1 intracellular trafficking itinerary.

The VINE complex could sort Mrl1 by recognizing sequences in its cytosolic tail or bind another protein that interacts with the MPR-like lumenal domain of Mrl1. A Vps10 mutant missing its cytoplasmic tail (Vps10^ΔCt^) lacks sorting signals and is transported to the vacuolar membrane and lumen (Bean et al., 2017; Cereghino et al., 1995; Cooper and Stevens, 1996). We hypothesized that if the Mrl1 tail contains a signal for VINE-mediated sorting, transplanting it onto Vps10^ΔCt^ (Vps10^ΔCt^-Mrl1^Ct^; Figure 6G) would confer VINE-dependent effects. We found that expression of *VRL1* did not alter the localization of WT Vps10 or Vps10^ΔCt^, the latter of which was targeted to the vacuolar membrane (Figure 6H). Vps10^ΔCt^-Mrl1^Ct^ localized to perivacuolar puncta and the vacuolar membrane in *vrl1* cells, indicating the presence of VINE-independent sorting signals in the Mrl1 tail (Figure 6H). *VRL1* expression caused a significant decrease in the proportion of cells displaying punctate Vps10^ΔCt^-Mrl1^Ct^ (14% decrease relative to *vrl1*, P < 0.001; Figure 6I), suggesting that the Mrl1 cytoplasmic tail is sufficient to confer VINE complex-mediated transport.

Because VINE redistributes Mrl1 from endosomes to the vacuolar membrane, we hypothesized that Mrl1 follows a Golgi-vacuole-endosome-Golgi recycling loop and that loss of Vrl1 specifically delays endosome to Golgi retrograde transport. Atg27, which follows a similar itinerary, relies on Snx4 to recycle it from vacuoles to the endosome (Suzuki and Emr, 2018). We observed that Mrl1-mScI was mislocalized to the vacuolar membrane in a *snx4Δ* strain (Figure 6J), which caused a significant decrease in endosomal Mrl1-mScI (P < 0.0001; Figure 6K), suggesting that Mrl1 is partially transported from the vacuole to the endosome by Snx4-containing coat complexes. Taken together, these results support a model where VINE enhances the recycling of the cargo protein Mrl1 at endosomes, causing a change in its steady state distribution (Figure 6L).

## Discussion

We have identified a novel endosomal SNX-BAR complex composed of the VPS9-domain GEF Vrl1 and the Vps5 paralog Vin1 which we have named the VINE complex. Our work suggests that VINE forms a novel class of endosomal transport carriers and sorts a unique set of cargo proteins that includes the mannose 6-phophate receptor-like protein Mrl1.

### Divergent N-terminal sequences in paralogous SNX-BARs specify complex formation

The function of the Vps5-related SNX-BAR protein Vin1 was not previously known. We found that in the absence of Vrl1, Vin1 is unstable and displaced to the cytosol suggesting that it functions solely as a member of the VINE complex. Vin1 and Vrl1 are both predicted to have PX-BAR domains and dimerize through a canonical BAR-BAR interface, yet this interaction is not the primary driver of Vrl1-Vin1 association. Instead, we found that a short sequence within the unstructured N-terminal extension of Vin1 binds specifically to the AnkRD of Vrl1, and this is necessary for selective incorporation into the VINE complex. Our work suggests that VINE assembly requires two inputs: a strong interaction involving the Vin1 N-terminus and a weak interaction between the BAR domains that may occur primarily at the endosomal membrane.

The Vin1 paralog Vps5 also has an unstructured N-terminus that is critical for its assembly with the retromer complex and binds to a conserved patch on the Vps29 subunit (Collins et al., 2005; Seaman and Williams, 2002). Moreover, the interaction between Vps5 and Vps17 PX-BAR domains requires chemical crosslinkers to detect in detergent-solubilized lysates (Horazdovsky et al., 1997), suggesting the interaction is weak or mediated largely by hydrophobic interactions. This supports a model for retromer assembly that parallels that of the VINE complex, where the Vps5 N-terminus makes critical interactions with other retromer subunits and drives the assembly of the Vps5-Vps17 BAR-BAR dimer through an avidity effect.

Vin1 and Vps5 arose from a single ancestral gene during a whole-genome duplication (WGD) event (Byrne and Wolfe, 2005; Wolfe and Shields, 1997), and subsequently diverged to assume new roles in the VINE and retromer complexes, respectively. In pre-WGD species, the single ancestral form of Vps5 must partner with both Vrl1 and Vps17, which requires some promiscuity in BAR-BAR pairing. This is consistent with our ab initio modeling, which predicted a variety of pairings between the PX-BAR domains of Vps5, Vps17, Vrl1 and Vin1, while the PX-BAR protein Mvp1 was predicted to form only homodimers, consistent with in vivo observations (Suzuki et al., 2021).

Van Weering et al. (2012) have proposed a lock and key model to explain the specificity of SNX-BAR pairing. In this model, paired charges within the hydrophobic BAR-BAR interface enforce the specificity of BAR-BAR interactions, and loss of these charged residues results in more promiscuous BAR pairing. Promiscuous BAR-BAR coupling could explain why interactions mediated by the N-termini of Vin1 and Vps5 are needed to specify complex assembly.

New functions have been uncovered for the extended N-termini of other SNX-BAR proteins, suggesting these extended regions have previously unappreciated regulatory roles. SNX1, which is the human homolog of Vps5 and Vin1, recognizes SNX5 (or its homolog SNX6) through BAR-BAR interactions based on lock-and-key charge pairing, and engages with other complexes, including SNX27 (Simonetti et al., 2021; Yong et al., 2021) and the retromer subunit VPS29 (Swarbrick et al., 2011), through its unstructured N-terminal domain. Thus, the N-termini of the SNX1/Vps5/Vin1 family of SNX-BAR proteins have diversified to bind different proteins and participate in different sorting complexes.

### The VINE complex is both a SNX-BAR coat and a VPS9-domain GEF

The VINE complex is the first described SNX-BAR coat to possess a VPS9 domain-containing subunit. Retromer binds the VPS9-domain GEFs Vps9 and Muk1, which redundantly activate Rab5-like GTPases to stimulate PI3P production at endosomes (Bean et al., 2015). One benefit of wiring SNX-BARs to VPS9-domain GEFs could be to generate a local enrichment of PI3P that enhances SNX-BAR assembly. Indeed, we find that VINE localization requires its own GEF activity in a strain lacking other VPS9-domain GEFs.

The human Vrl1 homolog VARP also contains a VPS9 domain and associates with retromer (Hesketh et al., 2014), albeit through a distinct interaction involving a motif that is not present in Vrl1 (Crawley-Snowdon et al., 2020). This example of convergent evolution suggests that the linking of retromer to the activation of endosomal Rabs has an important and conserved role. VARP activates Rab21, which is related to Rab5 (Stenmark and Olkkonen, 2001) and interacts with PI3K in a proximity-based assay (del Olmo et al., 2019), though it has not yet been shown to stimulate PI3K activity. Because endosomal Rab GTPases also recruit a variety of effectors including the conserved tethering complexes Rabenosyn-5/Vac1 and CORVET (Cabrera et al., 2013; Christoforidis et al., 1999; Peplowska et al., 2007; Peterson et al., 1999) further work is required to clarify the conserved functional link between VPS9-domain GEFs and SNX-BAR sorting complexes.

### VINE forms endosomal transport carriers and regulates cargo distribution

Our work suggests that VINE is a novel sorting complex that acts in a pathway-specific manner, thus joining the group of SNX-BAR complexes that promote independent sorting pathways from the endosome or vacuole in yeast (Ma and Burd, 2020; Suzuki et al., 2021). We identified the MPR-related protein Mrl1 as a candidate VINE cargo. Although the function of Mrl1 is unclear, there is evidence that it works jointly with Vps10 to enhance the transport or maturation of some vacuolar proteases (Whyte and Munro, 2001). Restoring VINE function alters the steady state localization of Mrl1 and could modulate the rate at which it delivers proteins to the vacuole. The metabolic implications of VINE-mediated Mrl1 sorting and the extent to which Mrl1 uses redundant SNX-BAR trafficking pathways are important topics for future study.

VARP, like Vrl1, also has a cargo-sorting role, as it binds to VAMP7 and cooperates with retromer to direct it on an endosome to cell surface pathway (Crawley-Snowdon et al., 2020; Hesketh et al., 2014; Schäfer et al., 2012). Additional research will be needed to understand the unique functions of the VINE complex in protein sorting and clarify its functional conservation with VARP. Importantly, almost all studies on endosomal trafficking and signaling have been performed in strains that lack the VINE complex, and we anticipate that many other VINE cargos exist. Some proteins that are recycled by SNX-BAR-dependent pathways, such as phospholipid flippases, are themselves important for organelle function or signaling (Dalton et al., 2017; Liu et al., 2008). In the absence of functional VINE, such proteins may be missorted and/or have altered properties. Restoring VINE may therefore restore cellular processes and reveal new biology.

## Materials and Methods

### Yeast strains and plasmids

Yeast strains and plasmids used in this study are described in Tables S4 and S5, respectively. Yeast strains were built in the BY4741 strain background using homologous recombination-based integration unless otherwise indicated. Gene deletions, promoter exchanges and tags were confirmed by colony PCR and either western blot or fluorescence microscopy where possible. Plasmids were built by homologous recombination in yeast, recovered in *Escherichia coli* and confirmed by sequencing.

### Bioinformatic analysis of protein folding and sequence conservation

Prediction of protein structure and binding interfaces was performed using Phyre2 (Kelley et al., 2015) and the ColabFold advanced server (Mirdita et al., 2021) with default settings. Vrl1 amino acid sequences were from the *S. cerevisiae* strain RM11-1a. Orthologous sequences were obtained from the OrthoDB database (Kriventseva et al., 2019), aligned using the EMBL-EBI Multiple Sequence Comparison by Log-Expectation (MUSCLE) tool (https://www.ebi.ac.uk/Tools/msa/muscle) and presented using Jalview (http://www.jalview.org).

### DHFR protein fragment complementation assay and ontology analysis

A MATa strain containing a plasmid that expresses the Vrl1-DHFR[1,2] (DHFR^Nt^) fusion from the *ADH1* promoter, or the p*ADHpr*-*VRL1(1-465)-DHFR^Nt^* control, was crossed into a library of MATα strains (n = ~ 4,300) expressing proteins fused to DHFR[3] (DHFR^Ct^; Tarassov et al., 2008). Diploids were subjected to two rounds of double mutant selection followed by two rounds of selection on media containing 200 µg/ml methotrexate, in 1536 arrays. Manipulations were carried out using a BM3-BC pinning robot (S&P Robotics inc., Toronto, Canada). Colony area was analyzed using CellProfiler (Lamprecht et al., 2007) after 8 days at 30 °C. Z-scores were generated using median colony area from two technical replicates for each Vrl1-prey combination. Functional analysis of Vrl1 DHFR interactors (Z > 2) was performed using the Gene Ontology (Ashburner et al., 2000; Gene Ontology Consortium, 2021) GO Enrichment Analysis tool (Mi et al., 2019).

### Fluorescence microscopy and automated image analysis

Yeast cells were diluted from overnight cultures in fresh synthetic dextrose-based media (SD) and incubated at 30 °C for ~4 hours or until they reached an optical density of ~ 0.4 - 0.7 OD_600_ unless otherwise indicated. Log phase yeast were transferred to concanavalin A-treated 96-well glass bottom plates (Eppendorf, Hamburg, Germany) and imaged using a DMi8 microscope (Leica Microsystems, Wetzlar, Germany) equipped with an ORCA-flash 4.0 digital camera (Hamamatsu Photonics, Shizuoka, Japan) and a high-contrast Plan-Apochromat 63x/1.30 Glyc CORR CS oil immersion lens (Leica Microsystems, Wetzlar, Germany). Image acquisition and processing was performed using the MetaMorph 7.8 software package (MDS Analytical Technologies, Sunnyvale, California). Yeast vacuoles were labelled with 100 µM CMAC (Setareh, San Jose, California) or 4 µM FM4-64 (Life Technologies, Carlsbad, California) for 30 minutes at 30 °C. Dye-treated cells were washed once in SD media prior to imaging.

Linear intensity scale changes were uniformly applied to all images in an experimental set using MetaMorph 7.8 (MDS Analytical Technologies, Sunnyvale, California). Images were prepared for presentation using Photoshop CC 2020 (Adobe, San Jose, California) and Illustrator CC 2020 (Adobe, San Jose, California). Quantification was performed on unscaled raw imaged with scripted MetaMorph 7.8 journals (MDS Analytical Technologies, Sunnyvale, California). The Count Nuclei feature was used to filter out dead cells and identify live cells based on intensity about local background (IALB). The Granularity feature was used to identify puncta in a dead cell-masked intermediate image based on IALB. Masking functions were performed using the Arithmetic function with Logical AND.

### Coimmunoprecipitation, western blotting and spheroplasting

For western blot-based stability assays, yeast cells were grown to log phase in SD media at 30 °C and 10 OD_600_/ml equivalents of cells were harvested and stored at −80 °C. Cells were thawed and lysed by vortexing in 100 µl of Thorner buffer (8 M Urea, 5% SDS, 40mM Tris-Cl (pH 6.4), 1% beta-mercaptoethanol and 0.4 mg/ml bromophenol blue) with ~100 µl of acid-washed glass beads/sample at 70 °C for 5 minutes. Lysates were centrifuged at 14,000 RPM for 30 seconds and separated on 8% SDS-PAGE gels followed by western blotting with mouse anti-HA (H9658, Clone HA-7, MilliporeSigma) or anti-PGK1 monoclonal antibodies (AB_2532235, 22C5D8, Invitrogen), and secondary polyclonal goat anti-mouse antibodies conjugated to horseradish peroxidase (115–035-146; Jackson ImmunoResearch Laboratories).

For CoIPs, yeast cells were grown to log phase in SD media at 30 °C and 75 OD_600_/ml equivalents of cells were incubated in 50 mM Tris-Cl with 10 mM DTT (pH 9.5) for 15 minutes at room temperature and digested in spheroplasting buffer (1.2 M sorbitol, 50 mM KH_2_PO_4_, 1 mM MgCl_2_ and 250 µg/ml zymolase at pH 7.4) at 30 °C for 1 hour. Spheroplasts were washed twice with 1.2 M sorbitol, frozen at −80 °C, then incubated in 500 µl of lysis buffer (0.1% Tween-20, 50 mM HEPES, 1 mM EDTA, 50 mM NaCl, 1 mM PMSF and 1x fungal ProteaseArrest, pH 7.4) at room temperature for 10 minutes. 50 µl volumes of lysate were collected for each sample and mixed with 2x Laemmli buffer (4% SDS, 20% glycerol, 120 mM Tris-Cl (pH 6.8), 0.01g bromophenol blue and 10% beta-mercaptoethanol) for western analysis while remaining lysates were incubated with either a polyclonal rabbit anti-GFP (EU2, Eusera) or a polyclonal rabbit anti-HA antibody (ab9110, Abcam) at 4 °C for 1 hour. Antibody-treated samples were next incubated with Protein A-Sepharose beads (GE Healthcare) at 4 °C for 1 hour. Beads were washed 3x in lysis buffer before being resuspended in 50 µl of Thorner buffer and heated at 80 °C for 5 minutes. Western blotting of proteins separated on 8% SDS-PAGE gels was carried out with monoclonal mouse anti-HA (H9658, Clone HA-7, MilliporeSigma), monoclonal mouse anti-HA (MMS-101R; Covance) or monoclonal mouse anti-GFP antibodies (11–814–460-001; Roche) prior to secondary antibody treatment with polyclonal goat anti-mouse conjugated to horseradish peroxidase (115–035-146; Jackson ImmunoResearch Laboratories). Blots were developed with Amersham ECL (RPN2232, Cytiva) or Amersham ECL Prime (RPN2236, Cytiva) chemiluminescent western blot detection reagents and exposed using Amersham Hyperfilm ECL (GE Healthcare). Densitometry of scanned films was performed using ImageJ (Schneider et al., 2012).

### Correction of genomic vrl1 mutation using CRISPR-Cas9

A plasmid containing the Cas9 enzyme and a single guide RNA (sgRNA) targeting the *VRL1*-disrupting *yml003w* mutation was pre-cloned using small fragment golden gate assembly (Marillonnet and Grützner, 2020). BY4741 was co-transformed with linearized split *URA3* marker Cas9-sgRNA(*VRL1*) plasmid and PCR product containing the corrected *VRL1* sequence. Ura^+^ colonies were sequenced and an isolate with intact *VRL1* was frozen down and used for experiments.

### Statistical analysis of quantitative data

Statistical tests were performed using GraphPad Prism 9.1.0 (GraphPad Software, San. Diego, California) as indicated in figure legends with the appropriate post-hoc tests. Normality of data was assumed but not formally tested and hypotheses were measured against a threshold of 95% confidence (or P < 0.05). Graphs were made in Microsoft Excel 2019 (Microsoft, Redmond, Washington). Column charts represent the average value from biological replicates while scatter points represent data from individual replicates and are coloured by replicate. Error bars report the standard error of the mean value.

## Acknowledgements

We thank Dr. Christian Landry (Laval University, Quebec City, Canada) and Dr. Maya Schuldiner (Weizmann Institute of Science, Rehovot, Israel) for sharing yeast libraries, Drs. Bjorn Bean and Vincent Martin (Concordia University, Montreal, Canada) for sharing yeast CRISPR reagents and Dr. Luc Berthiaume (University of Alberta, Edmonton, Canada) for generously sharing rabbit anti-GFP serum.

We gratefully acknowledge funding support from the Natural Sciences and Engineering Research Council of Canada (grant 2016-04290 to EC and PGS-D3 Doctoral Scholarship to SPS); Canada Foundation for Innovation (Leading Edge Fund 30636); Canadian Institutes of Health Research (grants 247169 and 365914 to EC, CGS-M Frederick Banting and Charles Best Canada Graduate Scholarship to SPS and MSF); BC Children’s Hospital Research Institute Jan M. Friedman Graduate Studentship to SPS; University of British Columbia Catalyst Paper Corporation Affiliated Fellowship and 4-Year Doctoral Fellowship to SPS and University of British Columbia Medical Genetics Rotation Award to MSF.

## Author Contributions

Conceptualization, EC; Methodology, SPS, MSF and EC; Validation, SPS and MSF; Formal analysis, SPS and MSF; Investigation, SPS, MSF and MD; Resources, SPS, MSF and MD; Data curation, SPS and MSF; Writing – original draft, SPS and MSF; Writing – review & editing, SPS, MSF and EC; Visualization, SPS and MSF; Supervision, SPS and EC; Project administration, EC; Funding acquisition, EC.

## Supplemental Figure and Table Legends

**Figure 1 S1.**
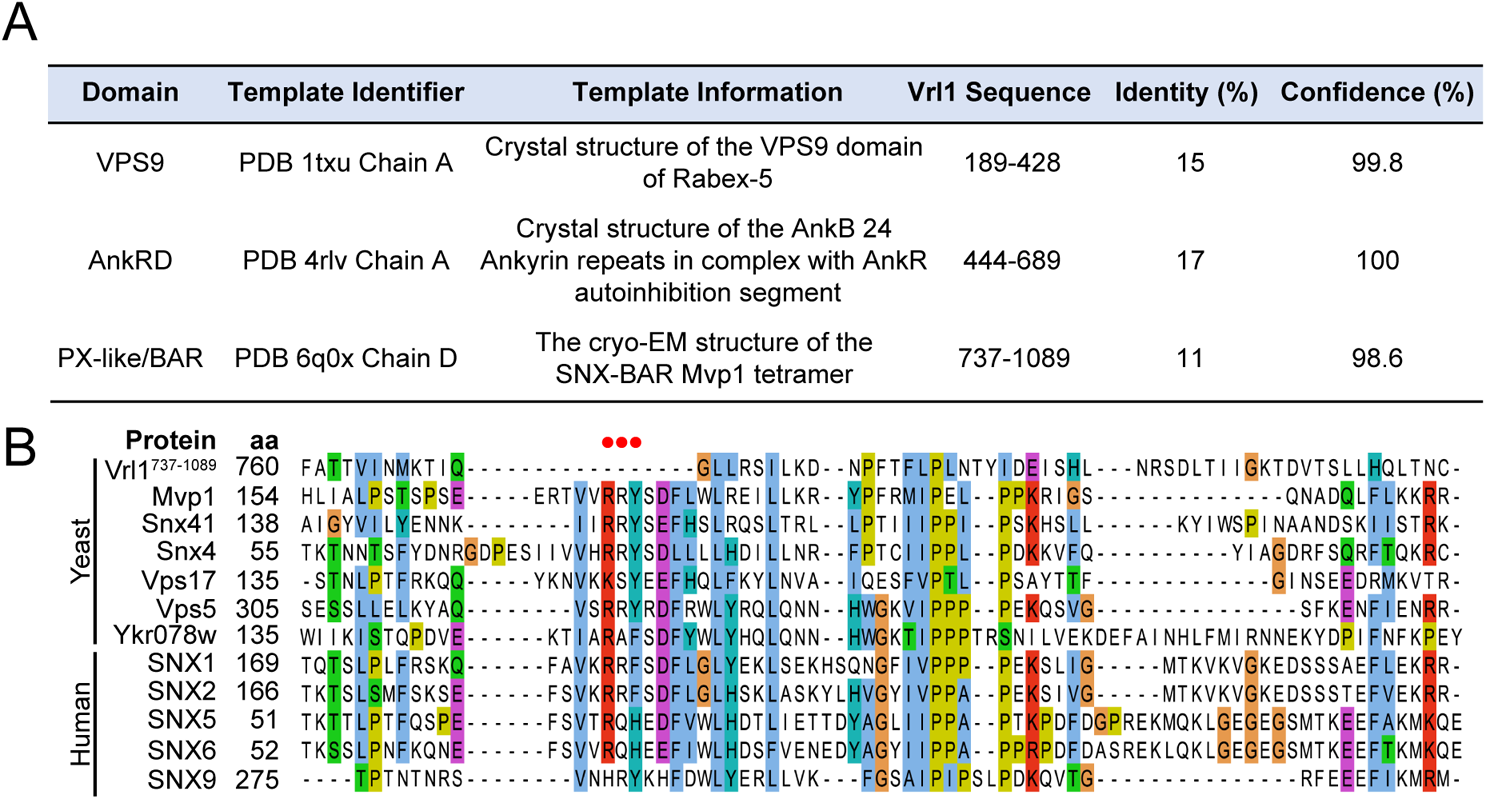
The Vrl1 PX-like domain is missing key PI3P-binding residues. (A) Results from Phyre2 analysis of Vrl1 sequences (Intensive mode, http://www.sbg.bio.ic.ac.uk/phyre2). (B) Sequence alignment of Vrl1 C-terminus (aa 737-1089) with yeast and human PX domain-containing proteins. Canonical PI3P-binding “RRY” motif is absent in Vrl1 (highlighted with red dots above alignment).

**Figure 1 S2.**
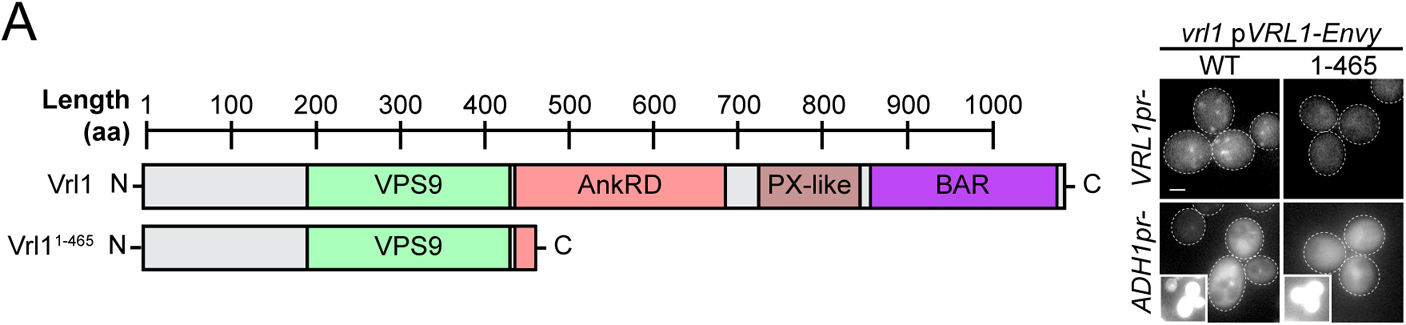
The Vrl1 N-terminus and VPS9 domain do not localize. (A) The Vrl1 N-terminus (aa 1-465) does not localize to puncta when expressed from the endogenous *VRL1* promoter or the strong *ADH1* promoter and instead accumulates in the cytosol. Scale bars, 2 µm.

**Figure 2 S1.**
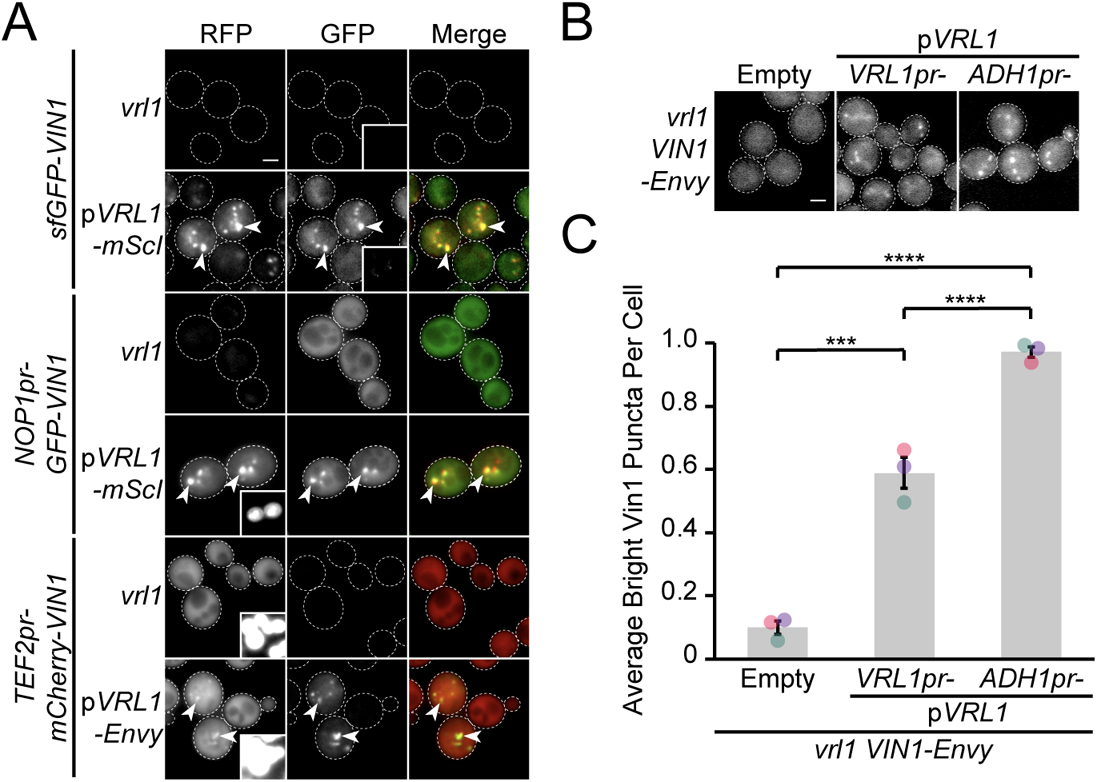
Vrl1 is indispensable for Vin1 puncta localization. (A) N-terminally tagged Vin1 requires Vrl1 for localization to puncta at three different expression levels. Insets are scaled to match other regions in the same channel. (B) Overexpression of Vrl1 from the *ADH1* promoter increases the average number of bright Vin1-Envy puncta per cell. (C) Quantification of Vrl1-Envy puncta per cell. One-way ANOVA with Tukey’s multiple comparison test; *n* = 3, cells/strain/replicate ≥ 1,495; *** = P < 0.001, **** = P < 0.0001. Scale bars, 2 µm. Error bars report SEM.

**Figure 3 S1.**
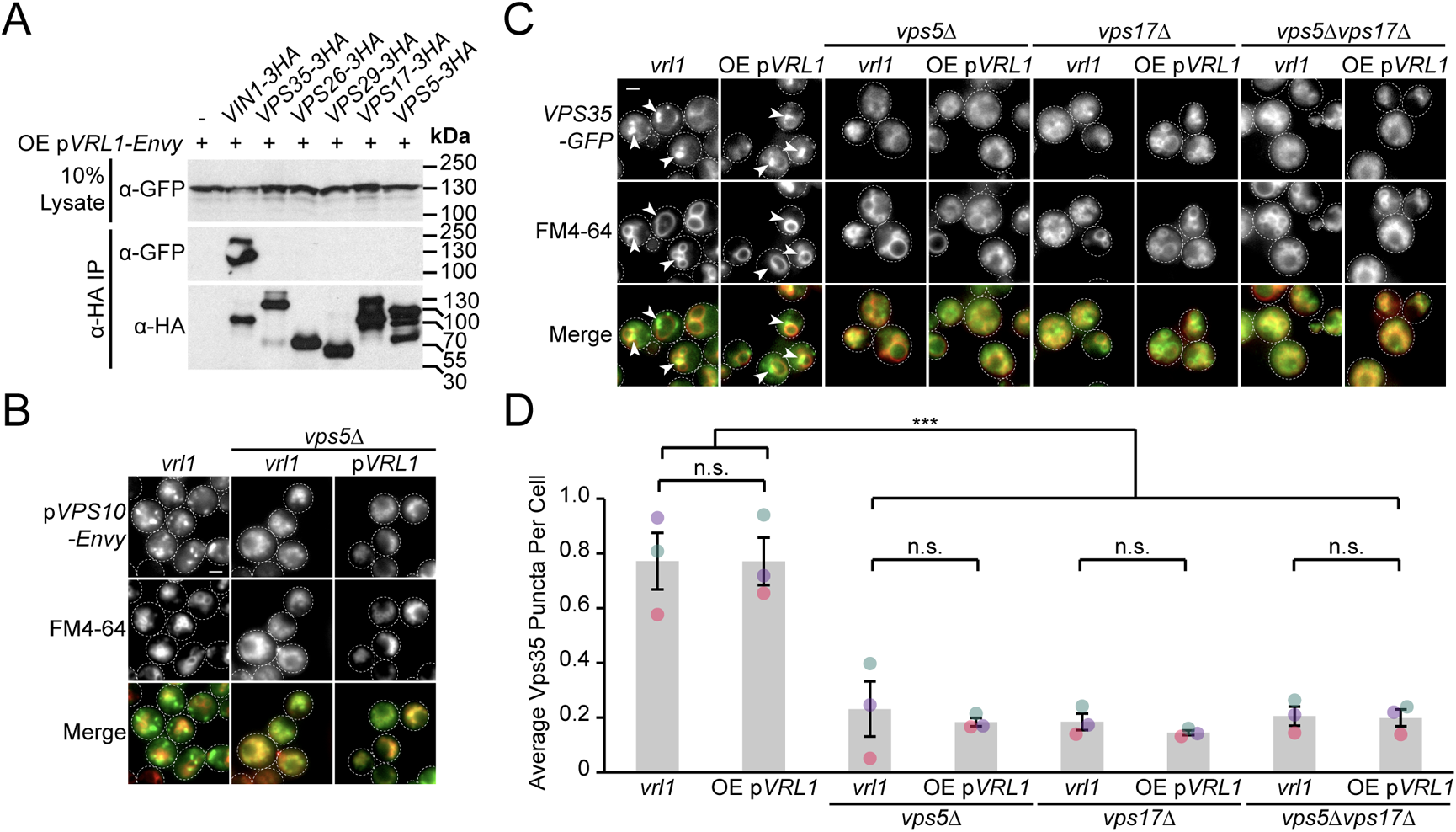
Vrl1 does not form a novel retromer-like complex. (A) Vrl1 does not interact strongly with subunits of retromer by CoIP. (B) Expression of *VRL1* does not rescue loss of endosomal Vps10-Envy in a *vps5Δ* strain. (C) Expression of *VRL1* does not rescue loss of endosomal Vps35-GFP or vacuolar morphology defects in retromer SNX-BAR deletion strains. (D) Quantification of Vps35-GFP puncta per cell. One-way ANOVA with Tukey’s multiple comparison test; *n* = 3, cells/strain/replicate ≥ 1,243; not significant, n.s. = P > 0.05, *** = P < 0.001. Scale bars, 2 µm. Error bars report SEM. OE, overexpressed.

**Figure 4 S1.**
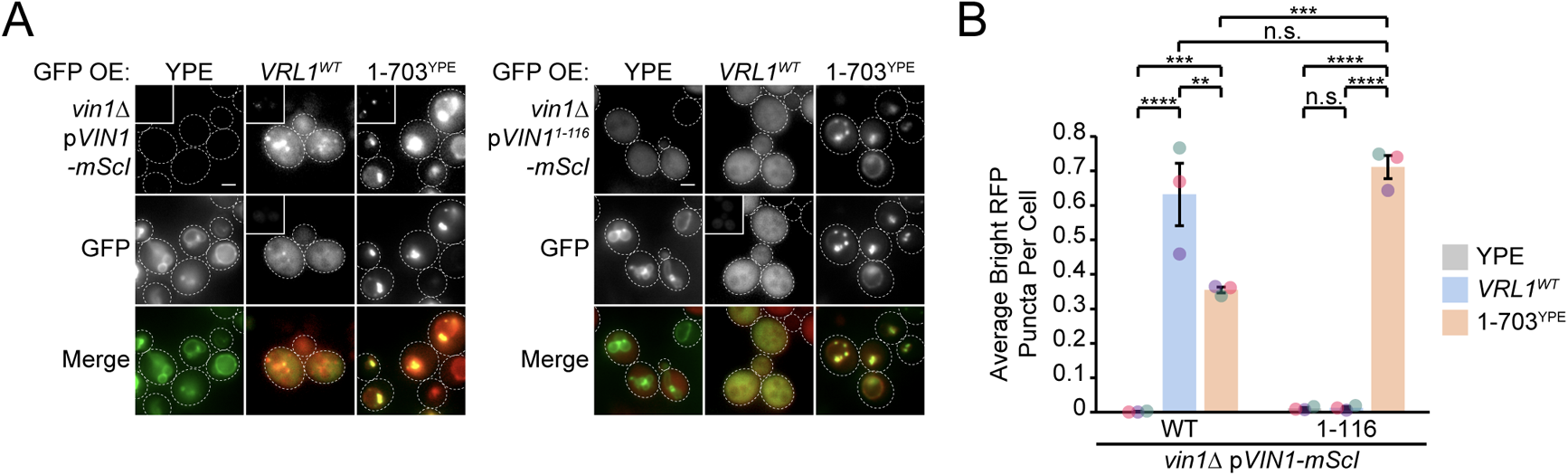
The Vin1 PX-BAR region is indispensable for Vrl1 localization. (A) The Vin1 N-terminus is not sufficient to localize overexpressed Vrl1-Envy to membranes in a *VIN1* deletion strain. The Vrl1-YPE chimera recruits the Vin1 N-terminal prey construct to puncta in strains lacking the chromosomal copy of *VIN1*. (B) Quantification of RFP puncta per cell. One-way ANOVA with Tukey’s multiple comparison test; *n* = 3, cells/strain/replicate ≥ 1,068; not significant, n.s. = P > 0.05, ** = P < 0.01, *** = P < 0.001, **** = P < 0.0001. Scale bars, 2 µm. Error bars report SEM.

**Figure 4 S2.**
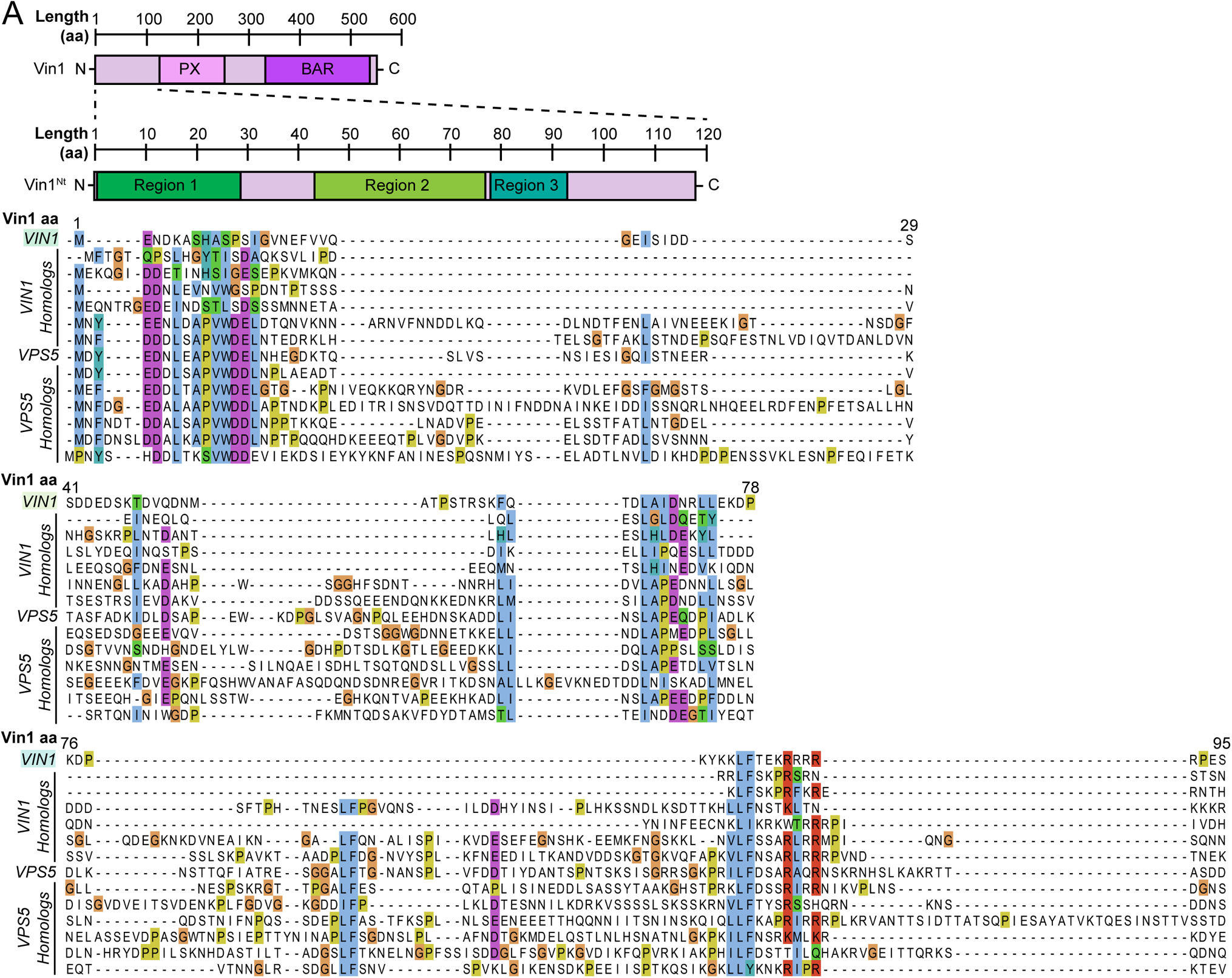
The Vin1 N-terminus has three conserved regions in fungal homologs. (A) Sequence alignment of Vin1 and Vps5 ohnologs collected using the Yeast Gene Order Browser (http://ygob.ucd.ie/). Three conserved regions were selected for expression as mScI-tagged fragments: Vin1^1-29^ (Region 1), Vin1^41-78^ (Region 2) and Vin1^76-95^ (Region 3).

**Figure 6 S1.**
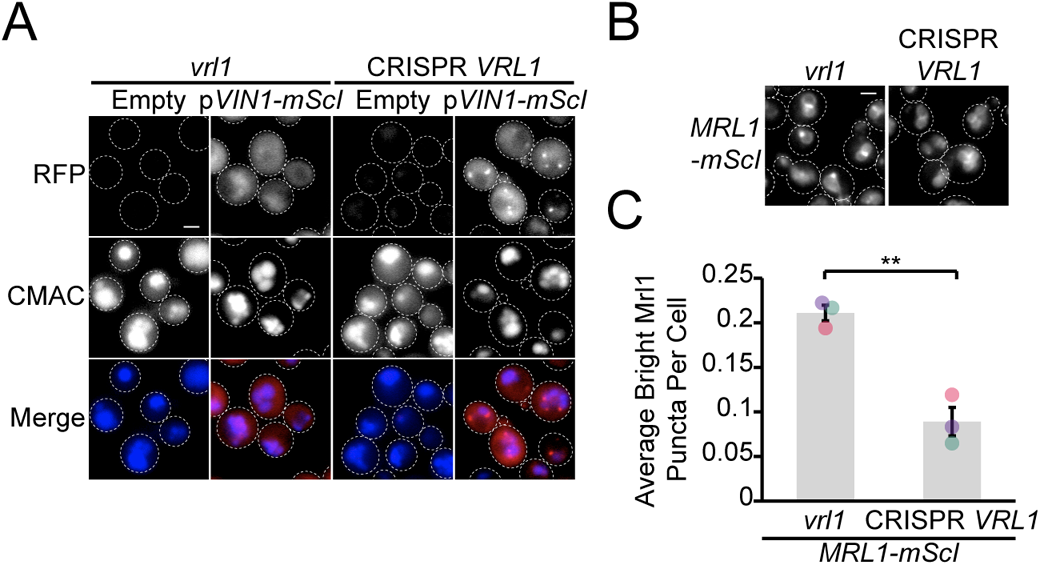
CRISPR-corrected Vrl1 recruits Vin1 and sorts Mrl1. (A) Localization of Vin1-mScI in BY4741 (*vrl1*) and an isogenic strain with the *vrl1* mutation corrected at the endogenous *VRL1* locus using CRISPR-Cas9 gene editing technology. Vin1-mScI localizes to puncta in a CRISPR-corrected *VRL1* strain. (B) Mrl1-mScI is redistributed from puncta in the CRISPR-corrected *VRL1* strain. (C) Quantification of large, bright Mrl1-mScI puncta per cell. Two tailed equal variance *t* test; *n* = 3, cells/strain/replicate ≥ 1,183; ** = P < 0.01. Scale bars, 2 µm. Error bars report SEM.

---

**Table S1. Vrl1 DHFR Interactors.** List of Z-scores from the Vrl1 DHFR screen.

**Table S2. Vrl1 DHFR Ontology Enrichment.** List of enriched ontology terms for Vrl1 DHFR interactors (Z > 2).

**Table S3. Yeast SNX-BAR Dimer Predictions.** Results of pairwise yeast SNX-BAR prediction matrix performed using ColabFold.

**Table S4. List of Saccharomyces cerevisiae strains used in this study.**

**Table S5. List of plasmids used in this study.**

